# Natural selection defines the cellular complexity

**DOI:** 10.1101/018069

**Authors:** Han Chen, Xionglei He

## Abstract

Current biology is perplexed by the lack of a theoretical framework for understanding the organization principles of the molecular system within a cell. Here we first studied growth rate, one of the seemingly most complex cellular traits, using functional data of yeast single-gene deletion mutants. We observed nearly one thousand *<u>e</u>*xpression *<u>i</u>*nformative *<u>g</u>*enes (EIGs) whose expression levels are linearly correlated to the trait within an unprecedentedly large functional space. A simple model considering six EIG-formed protein modules revealed a variety of novel mechanistic insights, and also explained ∼50% of the variance of cell growth rates measured by Bar-seq technique for over 400 yeast mutants (Pearson’s *R* = 0.69), a performance comparable to the microarray-based (*R* = 0.77) or colony-size-based (*R* = 0.66) experimental approach. We then applied the same strategy to 501 morphological traits of the yeast and achieved successes in most fitness-coupled traits each with hundreds of trait-specific EIGs. Surprisingly, there is no any EIG found for most fitness-uncoupled traits, indicating that they are controlled by super-complex epistases that allow no simple expression-trait correlation. Thus, EIGs are recruited exclusively by natural selection, which builds a rather simple functional architecture for fitness-coupled traits, and the endless complexity of a cell lies primarily in its fitness-uncoupled features.

## Introduction

Complex systems are widely observed in nature, and the embedded nonlinear relationships between components render the efforts for understanding a complex system often daunting and sometimes even beyond human capacity (*1*). A cell can be viewed as a complex system composed of a large number of genes each with dynamic expressions and together showing pervasive non-additive interactions (*2-6*). The central question of cell biology is to understand how genetic information flows in the complex molecular system to build up cellular phenotypes (*7*). Because analytical approaches are often unrealistic due to the lack of relevant theoretical frameworks, research strategies in cell biology are primarily empirical. Although substantial progresses have been achieved (*8*), it is unclear how much further such strategies can drive the field ahead because nonlinearity of the cell system compromises the practice of deduction (*9*). We reason that this issue can be gauged by the performance of explaining complex cellular phenotypes using experiences learned from previous data, and a good performance would suggest a tractable molecular system comprising a limited number of functional statuses in the cell.

## Results

The rate of cell growth is coordinated by nearly all aspects of biological processes and might be the most complex cellular trait. A variety of previous studies have attempted to model the cell growth of yeast *Saccharomyces cerevisiae* based on gene expression (*10-17*), but results are often lacking generality because of the limited functional space explored in each study; for instance, the expression of ribosome-associated genes can well predict the yeast growth rate across different environments (*13-15*), but is not correlated at all with the growth rate of natural yeast populations, whose variation is explained, instead, by the expression of amino acid biosynthesis genes (*17*). An ambitious reverse genetics project constructs a complete set of single-gene deletion mutants of *S. cerevisiae* (*18*), revealing that more than one third (∼2,000) of the yeast genes, when deleted, affect the cell growth rate by >5% in the rich medium YPD. With the recently available gene expression profiles of more than one thousand single-gene deletion mutants grown in YPD (*19*), we attempted to uncover the general functional architecture of cell growth rate of the yeast. The >1,000 yeast mutants with available expression profiles were divided into two random sets, with 885 mutants in Set #1 and the rest 443 in Set #2. We computed for every gene the correlation of its expression level to cell growth rate using the Set #1 mutants. Because growth rate is presumably controlled by numerous genes that interact with each other in a complex fashion, individual genes whose expression level correlates linearly to cell growth rate should be rare. However, we found more than 900 significant genes under a stringent statistical cutoff of *q* = 0.001 (Methods). The large number of such *e*xpression *i*nformative *g*enes (EIGs) suggested a relatively simple functional architecture that directly regulates the trait. To understand how the >900 genes are assembled to influence the growth rate we studied their protein-protein interactions. We uncovered six densely-connected protein modules, all of which are responsible for critical biogenesis processes, including “maturation of SSU-rRNA” for module #1 (M1), “amino acid biosynthesis” for module #2 (M2), “translation” for module #3 (M3), “cellular respiration” for module #4 (M4), “ribosomal large subunit biogenesis” for module #5 (M5), and “cell wall organization” for module #6 (M6) (table S1). We computed for each module its expression distance (ED) between the wild-type yeast and a given mutant, and, to avoid potential over-fitting, examined using the Set #2 mutants how cell growth rate can be explained by the EDs (Methods). Interestingly, we found that a linear function integrating the six EDs can explain ∼50% of the growth rate variation of the 443 Set #2 mutants (Pearson’s *R* = 0.69, *p* < 10^-16^; Fig. 1A). Note that the cell growth rates considered here are measured using Bar-seq technique (*20*), which is believed more accurate than the microarray-based method (*21*) or colony-size-based method (*4*), both used previously for quantifying growth rates of the yeast mutants. As far as the 443 Set #2 mutants are concerned, the Pearson’s *R* is 0.77 between the microarray-based measures and the Bar-seq-based measures, and 0.63 between the colony-size-based measures and the Bar-seq-based measures (Fig. 1B,C). Thus, the linear model was comparable to the two conventional experimental approaches in estimating cell growth rate of the yeast.

**Fig. 1.**
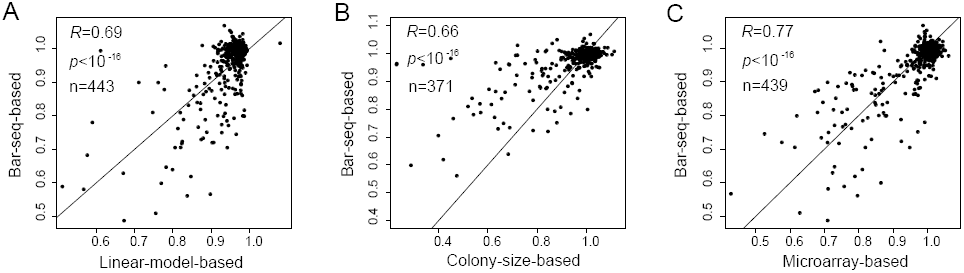
Comparison of the yeast cell growth rates estimated by the linear model or by the three experimental approaches. **(A)** The linear model is written as *G* = -1.740ED_*M1*_ - 0.435ED_*M2*_ - 0.725ED_*M3*_ - 0.071ED_*M4*_ + 0.794ED_*M5*_ - 0.058ED_*M6*_ + 1.019, where *G* stands for growth rate. **(B)** Seventy-two mutants without the colony-sized-based estimation are excluded. **(C)** Four mutants without the microarray-based estimation are excluded. Each dot represents a deletion mutant, with the Pearson’s *R* shown.

Among the six EIG modules M1 and M2 are formed primarily by essential genes, whereas the rest four mostly by non-essential genes, half of which, when deleted, show nearly normal growth (Fig. 2A). To find out whether there are major signal-contributing members, for a given module we removed each time a single gene and then checked its performance in explaining the cell growth variation. No matter whether they are essential or not, individual genes appeared to have a similar level of weak contribution to the overall performance of a whole module (Fig. 2B). We examined the module-level activities in mutants with a growth rate less than 80% of the wild-type. With only a few exceptions, the 87 slow-growth mutants form five clusters, each corresponding to the alterations of distinct modules (Fig. 2C). This pattern suggested that the six biogenesis-related modules represent rather independent causal factors of the growth defects, which helped clarify the previous confusion regarding the effects of ribosome-related genes (M5) and amino acid biosynthesis genes (M2) on the yeast cell growth (*17*). Note that we failed to observe such slow-growth mutant clusters based on the expressions of all individual genes of these modules (fig. S1). We conducted partial correlation analysis to reveal the potential epistasis between the modules. Strikingly, the Pearson’s *R* between ED_*M5*_ and the growth rate changed from -0.4 to 0.3 after controlling for the influences of the other modules (Fig. 2D), which can also be seen from the linear model used in Fig. 1A, in which the sign of ED_*M5*_ is positive. Because M5 represents ribosomal biogenesis that consumes up to 80% of the total cell energy (*22*) and its expression divergence (ED) is primarily due to the reduced gene expressions compared to the wild-type, it is likely that suppression of M5 *per se* saves energy, which promotes cell growth given alterations of the other modules often have already reduced the growth rate beneath a critical level. Consistently, deletion of *SSF1*, a member gene of M5, can be rescued by further deletion of *RPL16A*, a member gene of M3, or *PRM5*, a member gene of M6 (*4*) (Fig. 2E). This finding challenged the common belief that down-regulation of ribosomal genes reduces cell growth rate (*13-15*). These analyses collectively highlighted the strength of a general functional architecture in reconciling the diverse data and in revealing mechanistic insights.

**Fig. 2.**
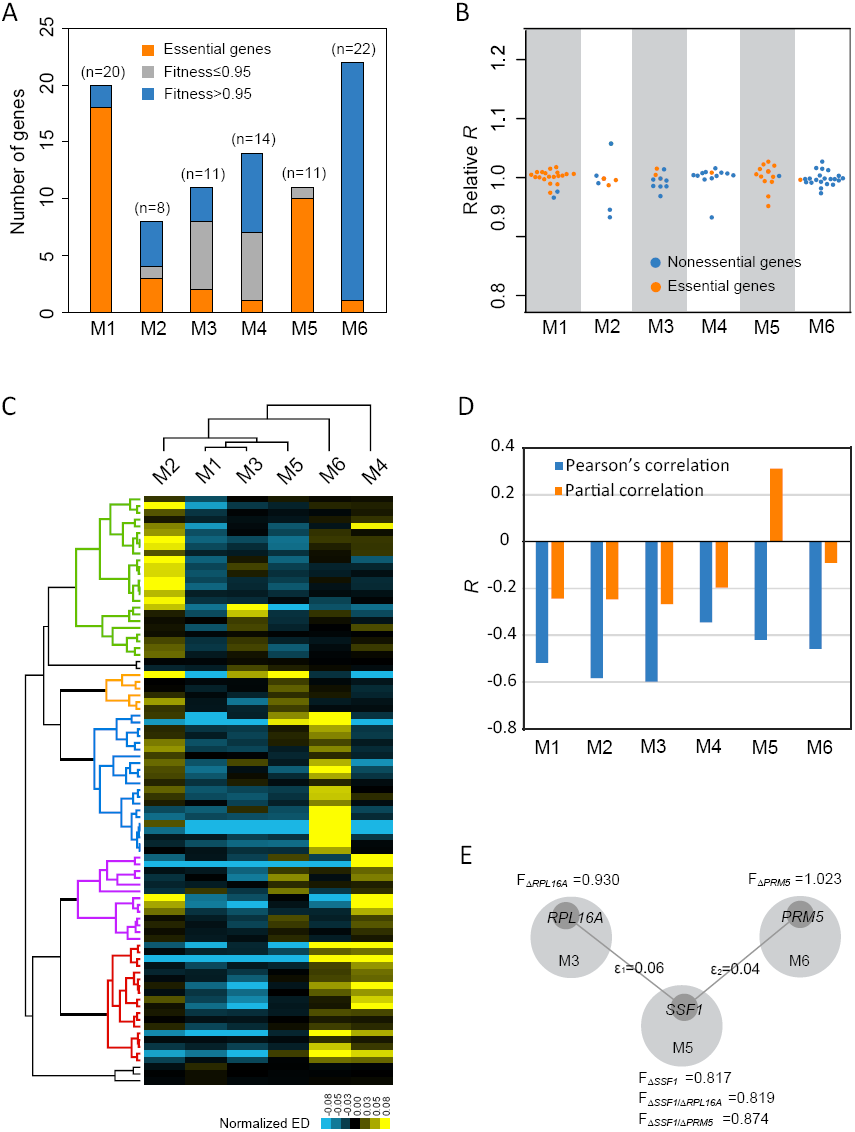
Characterization of the six growth-related EIG modules. **(A)** Composition of each of the six modules. **(B)** Effects after removing a single member gene on the correlation between module activity and cell growth rate for each of the six modules. The y-axis shows the Pearson’s *R* after removing a member gene relative to that of the whole module. **(C)** The five types of growth defects defined by the six modules. Each row represents a slow-growth mutant, and the expression distance (ED) of a module is normalized by subtracting its mean ED in the 87 mutants. **(D)** The Pearson’s *R* between module activity and cell growth rate for each of the six modules, in comparison to that of the partial correlation that controls for the other five modules. **(E)** The rescuing epistasis between *SSF1* of M5 and *PRL16A* of M3 or *PRM5* of M6. F represents the relative growth rate (or fitness) of a mutant, with ε_1_ = F_Δ*SSF1*/Δ*PRL16A*_ - F_Δ*SSF1*_ x F_Δ*PRL16A*_ and ε_2_ = F_Δ*SSF1*/Δ*PRM5*_ - F_Δ*SSF1*_ x F_Δ*PRM5*_.

By analyzing the microscopic images of triple-stained yeast cells a previous study characterized 501 morphological traits for the ∼5,000 yeast mutants each lacking a non-essential gene (*23*). We identified for each of the traits its EIGs using the same Set #1 mutants. Up to 2,541 non-redundant genes were identified as EIGs of at least one trait, and mean and median number of traits an EIG affects were 27 and 11, respectively. The number of EIGs a trait has varied substantially, ranging from zero to ∼1,000. Surprisingly, there was little overlap between EIGs and the corresponding *g*enetically *i*nformative *g*enes (GIGs) that, when deleted, result in significant effects on the traits (*24*) (fig. S2). Compared with EIGs, GIGs tend to regulate a much larger number of downstream genes but tend not to respond to other genes (Fig. 3A). It is thus likely that the significant effect on a trait requires perturbations to multiple EIGs of the functional modules that directly regulate the trait, which is achieved often by deleting a GIG with many downstream targets rather than a single EIG. This notion, if correct, would help clarify a long-standing puzzle that genes with expression response to a given environment are often not the genes required for the environment (*18*). Notably, although the diverse genetic perturbations can effectively remove coincidental associations between gene expression level and a trait, still some of the EIGs may regulate while the others may be reactive to their traits (*25, 26*). By examining the segregation pattern of trait, QTLs and gene expression in the F1 segregants of a hybrid of two *S. cerevisiae* strains with distinct genetic backgrounds (*27*) (Methods), we estimated that approximately 15-40% of the EIGs can causally affect the corresponding morphological traits (Fig. 3B). We further dissected 10 exemplar traits each with >300 EIGs, and found that protein modules formed by these EIGs have generally good performance in explaining each of the traits, with Pearson’s *R* ranging from 0.3 to 0.6 (fig. S3 and table S2).

**Fig. 3.**
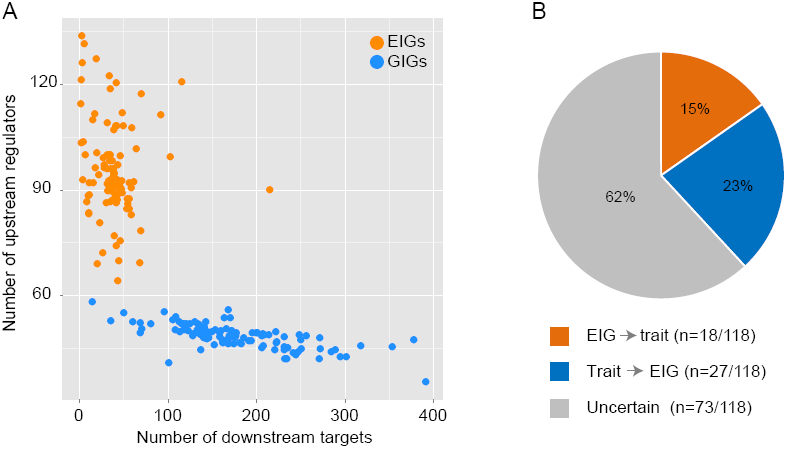
Functional characterization of GIGs and EIGs. **(A)** The numbers of downstream targets (x-axis) and upstream regulators (y-axis) per EIG or GIG. Each dot represents the average of all EIGs or GIGs of a trait. **(B)** Assessment of the causality for the 118 EIG-trait associations, with 18 EIG -> trait and 27 trait -> EIG causal relationships determined under a false discovery rate of 0.05.

However, virtually no EIGs were found for a large number of traits that have as complex genetic architectures as others; for example, there were on average 129 ± 22.9 (mean ± s.e.m.) GIGs for the 48 traits with <10 EIGs, and this number was 167 ± 15.6 (mean ± s.e.m.) for the rest 168 traits with ≥ 10 EIGs (*p* > 0.05, Mann-Whitney U test; Fig. 4A). We reasoned that their underlying functional architecture should be composed of complex gene-gene interactions that preclude the detection of EIGs, a process relying on the rather simple expression-trait relationship. Interestingly, the relatedness of a morphological trait to cell growth rate appeared to be the determinant of the EIG number. Specifically, although there were typically several hundred EIGs in traits tightly coupled with cell growth rate, we found no EIGs in traits with no significant correlation to the cell growth rate (Fig. 4B and fig. S4). This finding cannot be explained by the noise of trait measuring (fig. S5) or by a smaller variation of the traits less coupled with cell growth rate (fig. S6). In addition, we here required the EIGs found in the Set #1 mutants reproducible in the Set #2 to avoid false positives. Because for the single-celled yeast cell growth rate represents the organism’s fitness, the above pattern indicated that natural selection has been critical in shaping the composition of EIGs. Specifically, natural selection seemed to be necessary for the recruitment or maintenance of EIGs, and the absence of EIGs suggested a complex functional architecture underlying the fitness-uncoupled traits, as a result of neutral genetic drift or hitchhiking effects. On the other hand, given the generally good performance of the EIG-based functional architecture, it is likely that genes with a rather simple expression-trait relationship are preferentially recruited and maintained by natural selection, resulting in a much simpler functional architecture than anticipated for the fitness-coupled traits.

**Fig. 4.**
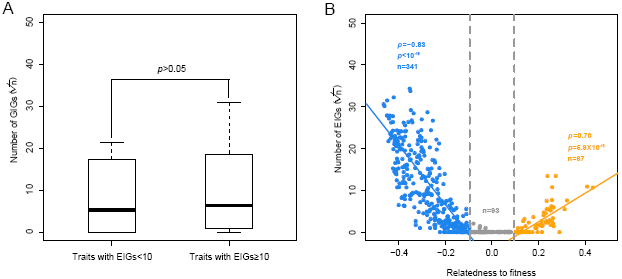
The presence of EIGs is determined by fitness coupling rather than genetic complexity. **(A)**The number of GIGs is comparable between traits with <10 EIGs (n = 48) and traits with more EIGs (n = 168). Box-plots are presented, and the y-axis shows the square root of the number of GIGs. **(B)** The number of EIGs as a function of trait relatedness to fitness. Each dot represents a trait, and the y-axis shows the square root of the number of EIGs. The trait relatedness to fitness is defined by the Pearson’s *R* between the trait values and cell growth rates of the deletion mutants, with *R* > 0.1 or *R* < -0.1 regarded as statistically significant after controlling for multiple testing (*q* < 0.01).

## Discussion

There are three caveats that warrant discussion. First, cell growth rate measured in YPD may not well represent the natural fitness of yeast, although the relative growth rates of the deletion mutants measured in diverse media are largely conserved (*21*), which may confound the estimations of fitness coupling of the morphological traits. This potential problem, however, is unlikely to generate the strikingly different EIG composition between the fitness-coupled and fitness-uncoupled traits; it would only erase the signature to make our analysis more conservative. Second, no gene-environment interaction was considered. We reasoned that, same as genetic factors, environmental factors affect traits also via modulating gene activities. The >1,300 single-gene deletion mutants represented diverse genetic perturbations (fig. S7), thus providing a reasonably good sampling of the entire functional space of the yeast cell, although only a single environment was examined here. In line with this, knowledge learned from the Set #1 mutants worked well in the independent Set #2 mutants. Third, because the genetic/phenotypic space represented by the F1 segregants of the BY x RM hybrid is limited, only a small subset of the identified EIGs can be tested for their causal effects on the traits. Although this gave a rough estimation of the proportion of causal EIGs, we still had to use all EIGs to build the functional architecture of the traits. Sampling more variations of the natural populations would help distinguish causal EIGs from reactive EIGs, complementing the present strategy that is based primarily on associations.

The genome-wide reverse genetic approach allowed us to explore an unprecedentedly large functional space of the yeast cell to identify an unbiased functional architecture of the yeast traits. Notably, the linear function considering the six biogenesis-related EIG modules were able to explain up to ∼50% of the variance of cell growth rate, one of the most complex traits, and also provided a variety of novel mechanistic insights on the regulation of yeast cell growth. This performance should be much improved if protein activity was considered in the context of a comprehensive interactome (*28*). In addition, the fact that ∼94% (82/87) of the slow-growth mutants can be assigned into five clear groups suggested that a more sophisticated model integrating the six modules may provide an even better understanding of the cell growth. Thus, there are good reasons to prospect a high-quality EIG-based functional architecture for most fitness-coupled cellular features, suggesting a manageable task for fully understanding them.

The most important finding of this study is probably the sheer absence of EIGs for most fitness-uncoupled traits, given the generally hundreds of EIGs observed for their fitness-coupled counterparts. Because these traits often have an as complex genetic architecture as the fitness-coupled ones (Fig. 3A) (*24*), the only explanation is that they are controlled by super-complex gene-gene interactions that erased all signatures of the expression-trait associations, rendering the task of revealing their functional bases at the gene level extremely difficult. How such complexity originated is intriguing. It may evolve specifically for regulating the traits via neutral genetic drift, or it may represent non-specific signals that are initiated by certain genetic perturbations and then propagating pervasively within the molecular system of a cell. Importantly, the lack of effective selection predicts that pathways mediating such regulations or signal propagations must be *ad hoc* settings with high turnover rate. This situation reminds us of the many known chaotic events observed in other complex systems that cannot be deterministically understood (*1*). Thus, the presence or absence of natural selection defines two types of biological phenomena, namely, the fitness-coupled and the fitness-uncoupled. Like linear and nonlinear differential equations in mathematics, where the former tends to have straightforward solutions but the latter is often unsolvable, the two kinds of biological phenomena should be studied and understood using different research philosophies and with different expectations. Recognition of this may save cell biology from the present endless complexity.

## Methods

### Data

The yeast *Saccharomyces cerevisiae* single-gene deletion stock was generated by Giaever et al. (2002), with 4,718 mutant strains each lacking a nonessential gene being considered in this study. As for cell growth rates of the above mutants measured in the rich medium YPD (yeast extract, peptone, and dextrose), the Bar-seq-based data were by Qian et al. (2012), the microarray-based by Steinmetz et al. (2002), and the colony-size-based by Costanzo et al. (2010). The 501 morphological traits of the mutants (SCMD) were characterized by Ohya et al. (2005), and the *<u>g</u>*enetically *<u>i</u>*nformative *<u>g</u>*enes (GIGs) that show significant phenotypic effects after deletion were defined for 220 traits by Ho and Zhang (2014), with 216 reproducible using the updated data in SCMD and thus included in this study. The microarray-based expression profiles of 1,484 deletion mutants were recently generated by Kemmeren et al. (2014), and gene A was called the downstream target of gene B and B was the upstream regulator of A if A shows significant expression change (*p* < 0.0001 as provided in the original data) in the mutant of B deletion.

### Identification of expression informative genes (EIGs)

There are 1,328 strains with both the expression profiles generated by Kemmeren et al. (2014) and the Bar-seq-based cell growth rates. We randomly divided the 1,328 yeast strains into two sets, with 885 strains for Set #1 and 443 for Set #2. There are 6,123 yeast genes on the chip used by Kemmeren et al. (2014). The distribution of cell growth rates of the mutants is highly biased, with the majority close to the rate of the wild-type. We thus computed for each of the 6,123 yeast genes the correlation of its expression levels with the corresponding growth rates using the univariate Cox’s regression model, with growth rate as the parameter “time”, strains of growth rate <0.9 weighted as “event = 1”, and all others as “event = 0”. Specifically, we first generated 500 artificial datasets, each containing 443 strains picked randomly from the 885 Set #1 strains with replacements. We then performed the Cox’s regression analysis for each of the 500 datasets, respectively, based on the settings described above, and obtained 500 *p*-values for every yeast gene. We defined the correlation robustness (*r*-value) of a given gene as the harmonic mean of its *p*-values after dropping both the highest and the lowest 5% of its 500 *p*-values, which was then multiplied by 6,123 for multiple testing correction. A total of 911 genes each with the corrected *r*-value<0.001 were defined as expression informative genes (EIGs) of the cell growth rate.

Different from the one-tailed distribution of cell growth rate, nearly all of the 501 morphological traits of the deletion mutants show a bell-shape distribution, with the median trait value very close to that of the wild-type (fig. S8). Thus, we used the Pearson’s linear regression model that considers the trait-gene correlation across the whole distribution, instead of the Cox’s regression model that emphasizes the difference of two categories. Specifically, we calculated the Pearson’s *R* for each of the 501 x 6123 trait-gene pairs in the 500 artificial datasets generated above. The Pearson’s *R* values were transformed into *p*-values using T-test, and the correlation robustness (*r*-value) was then computed as previously described. Genes with the corrected *r*-values less than 0.01 were considered as EIGs that are subject to the protein module analysis. To further reduce false positives, we required that these EIGs also show significant trait-gene correlation in the independent Set #2 mutants to be included in the analysis for Fig. 4 and it derivatives.

Morphological traits are not independent; for instance, the size and the diameter of a cell are correlated. To reduce correlated traits, we employed an un-supervised affinity propagation strategy proposed by Frey and Dueck (2007) to cluster the 501 traits based on the *r*-values of all genes, resulting 57 clusters each with an exemplar trait.

### Separation and functional annotation of EIG modules

The yeast protein-protein interactions (PPIs) were downloaded from BioGrid, a database built by Stark et al. (2006). For a given trait we constructed a non-directional, unweighted PPI network composed exclusively of its EIGs. Protein modules were separated using an order statistics local optimization method (OSLOM) proposed by Lancichinetti et al. (2011) with default settings. To annotate the biological functions of these protein modules, we performed the gene ontology (GO) enrichment analysis for each module using BinGO by Maere et al. (2005) and Cytoscape by Shannon et al. (2003). We obtained seven modules formed by the EIGs associated with cell growth rate, among which six were found to be enriched with functionally similar proteins under a false discovery rate of 0.001 (table S1). We identified 72 protein modules each with at least three genes for EIGs of the 10 exemplar morphological traits, and successfully annotated 66 using the GO enrichment analysis (table S2). Most of the annotated modules show >10-fold enrichment with the assigned function.

### Calculation of expression distance (ED)

For a given EIG module its expression distance (ED) between a mutant and the wild-type was defined as the Euclidian distance between the two expression profiles:

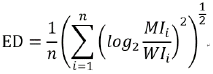

Where *MI*_i_ and *WI*_i_ are the expression level of the i^th^ gene in the mutant and wild-type strains, respectively, and n is the number of genes in the module.

### Determination of causal associations between the EIG expression and traits

Information of the genotype, expression and morphology of 62 F1 segregants of a hybrid of two yeast strains (BY4716, a derivative of S288c, and YEF1946, a derivative of RM11-1a) was obtained from Nogami et al. (2007), with three segregants excluded from further analyses because of unmatched IDs. Because there is no major difference between the two parental yeast strains in most of the morphological traits, there are only 118 EIGs whose expression-trait correlation was also detected in the 59 F1 segregants under *q* < 0.01 (two-tailed T test with Bonferroni correction for multiple testing). The causality of the EIG-expression versus trait association was resolved using the Network Edge Orienting (NEO) method developed by Aten et al. (2008). Following the manual provided by NEO, we calculated the LEO.NB.CPA score and the LEO.NB.OCA score with all genotype information (SNPs) inputted; for each association the two causality directions (i.e., EIG-expression -> trait and trait -> EIG-expression) were tested separately. We defined a cause relationship if the LEO.NB.CPA score > 0.8 and the LEO.NB.OCA score > 0.3, which corresponds to a false discovery rate of 0.05. We found 18 EIG-expression -> trait and 27 trait -> EIG-expression causal relationships. We were not able to assign a reliable causal relationship for the rest 118-18-27=73 associations.

### Calculation of the relatedness of the morphological traits to fitness

The relative cell growth rate is a reasonable measure of the relative fitness for the single-celled yeast. Because in this study all cellular traits are measured in YPD, we used the cell growth rate in YPD as the proxy of fitness.

Given the bell-shape distribution of a morphological trait where the wild-type trait value is almost always located in the middle, both increase and decrease of a trait value relative to the wild-type could affect fitness in the same direction. Thus, we divided for a given trait the 4,718 mutants into two equal halves according to the trait values, and calculated the Pearson’s *R* between trait value and fitness for each half of the mutants separately, resulting in two *R*s for every trait. The *R* with the larger absolute value was used to represent the relatedness of the trait to fitness. We also computed the Pearson’s *R* without separation of the mutants into two halves, and found that it is often highly similar to the relatedness obtained above (fig. S9).

There are 420 Set #2 mutants with the morphology information. Considering the bell-shape distribution of a morphological trait, we divided the 420 mutants into two halves, one with larger trait values and the other with smaller trait values, and identified the half that is more coupled with fitness. This half was used for testing the trait-specific EIG modules as shown in fig. S3.

### Assessment of the quality of the morphological traits

To characterize the yeast morphological traits Ohya et al. examined on average 400 individual cells for each mutant. The trait value of a given mutant is the mean trait value of the examined cells. Despite the generally large number of examined cells, for some traits there were often a few tens of informative cells, which may affect the reliability of the measurements. To address this issue, we randomly divided the examined cells of each mutant into two equal halves and computed the traits for each half separately. For each trait we then computed the Pearson’s *R* between values derived from the first half and values from the second half. The consistency between the two halves varies substantially among the traits, but is not dependent on the trait relatedness to fitness (fig. S5).

## Acknowledgements

We would like to thank Drs J. Zhang (U of Michigan), A. Dean (U of Minnesota), C-I Wu (U of Chicago), Y. Sheng (National U of Singapore), P. Shi (KIZ, CAS), W. Qian (IGDB, CAS), Y. Zhang (BIZ, CAS), S. Zhou (Baylor College of Medicine) for discussion or comments. This work is supported by two research grants from NSFC to X. H.. X. H. is supported by the Changjiang Scholars program and the Qing Nian Ba Jian program.

